# Drought-Spec-Net: Early Tomato Drought Detection and Potential Yield-Impact Assessment Using Vis–NIR Data

**DOI:** 10.64898/2026.09.24.754113

**Authors:** Kabir Hossain, Aniruddha Maiti, Karthik Chinnannan, Padma Nimmakayala, Eddie Sweeney, Zichun Wang, Natwange Chiwele, Luice Khamboo, Lakshmi Meghana Nallagari, Alexander Bucksch

## Abstract

Drought stress significantly reduces tomato (*Solanum lycopersicum* L.) productivity, and early detection is critical to minimize yield losses through timely interventions. In this study, we developed Drought-Spec-Net, a hybrid 1D convolutional neural network that integrates local and global spectral feature extraction to detect early drought stress from visible and near-infrared (Vis–NIR) spectra data of tomato seedlings. The model was trained on 378 samples using an 80:20 train–test split, with 20% of the training set reserved for validation. Drought-Spec-Net outperformed the evaluated baseline and state-of-the-art models, achieving 97% accuracy, 95% precision, 98% recall, and an F1-score of 97%. To improve the agronomic interpretation of the model outputs, predicted drought probabilities were converted into a literature-informed potential yield-impact indicator using a maximum impact level of 60%. On the test set (76 samples), mapped potential yield-impact values ranged from 0% to 60%, with an average reduction of 12.97%. We also conducted an initial experiment using our greenhouse RGB dataset, collected daily from drought-treated and well-watered tomato plants at West Virginia State University (WVSU). From this dataset, 44 images were selected for ilastik-based canopy segmentation, producing plant-level drought severity indices (DSI) with a mean of 0.28, median of 0.14, and range of 0.01–0.91. Additionally, we trained and fine-tuned a large language model (LLM) based on PLLaMA-7B-Instruct, called AgriLLaMA, for automated agronomic report generation from Drought-Spec-Net outputs. The generated reports summarize predicted stress levels, mapped potential yield impacts, and preliminary management considerations. This integrated approach not only improves early drought stress detection but also delivers quantitative and interpretable estimates of potential productivity losses, providing a complete framework connecting physiological stress detection to actionable agricultural outcomes.

## Introduction

With the global population projected to reach 9 billion by 2050, ensuring a stable food supply remains a critical concern^1,2^. Up to 40 percent of the world’s land area is already degraded, affecting nearly half of humanity^3^, further increasing pressure on agricultural systems. Ensuring global food security and supporting economic growth are closely tied to agricultural productivity, making efficient monitoring of plant health and crop conditions increasingly important^4^. Abiotic stress is a major constraint to agricultural productivity, with drought, heat, and waterlogging significantly affecting crop growth and yield^5^. Among these, drought stress is one of the most significant abiotic factors limiting agricultural productivity worldwide^6,7^, particularly in tomato^8^, where even short periods of water deficit can severely affect physiological processes such as photosynthesis^9^, nutrient transport^10^, and leaf water regulation^11^. These disruptions ultimately lead to substantial reductions in yield and quality^12^. Reported tomato yield losses under severe drought conditions can range from approximately 55% to 70%, depending on genotype, stress intensity, and environmental conditions^8,13^.

Beyond direct yield reduction, drought stress also affects fruit size, number, and marketability, making it a critical factor in both productivity and economic return. The relationship between drought stress and yield loss is complex and often nonlinear, as it depends on multiple interacting factors including drought intensity and duration, plant developmental stage, and adaptive physiological responses. Importantly, physiological responses to water deficit can develop before visible symptoms become apparent, creating a critical window for early stress detection and timely intervention before substantial productivity losses occur. Therefore, accurate early detection of drought stress is essential for enabling timely irrigation management and minimizing potential productivity losses^14^. Nevertheless, translating early stress signals into validated quantitative yield-loss predictions remains challenging, particularly when corresponding plant-level yield measurements are unavailable.

Traditional drought stress assessment methods rely on visual inspection or destructive physiological measurements, which are often time-consuming, subjective, and unsuitable for large-scale monitoring^15^. These approaches also limit continuous monitoring due to their invasive nature. In contrast, visible and near-infrared (Vis–NIR) spectroscopy provides a rapid and non-destructive alternative by capturing biochemical and structural changes in plant leaves associated with stress conditions^16^. These spectral signatures enable early discrimination between healthy and drought-stressed plants before visible symptoms appear^17^.

Recent advances in machine learning and deep learning have significantly improved the ability to analyze spectral data for plant stress detection^15^. Beyond spectral measurements, these approaches have demonstrated their value across different plant-sensing modalities. LeafyResNet used deep learning with UAV-acquired RGB imagery to detect Fusarium wilt and quantify disease severity in lettuce^18^. LeafyVGG-16 applied transfer learning to tomato disease and nutrient-deficiency classification while evaluating model vulnerability to adversarial attacks^19^. DIRTNet combined deep learning with in-soil fiber Bragg grating measurements for root-trait estimation and discrimination between control and drought conditions^20^. Focusing specifically on spectral drought-stress classification, previous studies have explored convolutional neural network (CNN)-based models^21^, residual learning frameworks, and classical machine-learning methods such as random forests (RFs)^22^ for drought-stress classification. More recently, specialized one-dimensional deep-learning architectures, including One-Dimensional Spectrogram Power Net (1D-SP-Net)^23^ and a one-dimensional CNN with an embedded residual global-context block (1D-ResGC-Net)^17^, have been developed for spectral-based drought-stress detection, achieving improved performance through enhanced feature representation and model design^17,23^.

Despite these advancements, most existing studies primarily focus on classification accuracy and stress identification, with limited attention to translating model predictions into agronomically meaningful outputs such as potential yield-impact assessment. While previous studies have examined drought-induced yield losses using field-based observations or statistical approaches, these approaches are often disconnected from real-time sensing and machine learning-based prediction systems. Furthermore, limited attention has been given to integrating model outputs into interpretable and user-oriented reporting frameworks, such as those enabled by recent advances in large language models^24^. As a result, a significant gap remains in linking early stress detection with quantitative yield impact assessment and actionable interpretation within a unified framework. Bridging this gap is essential for transforming predictive models into practical decision-support tools for farmers and agronomists.

To address these limitations, we propose Drought-Spec-Net, a hybrid one-dimensional convolutional neural network that integrates local convolutional feature extraction with global spectral representation learning through a parallel dense feature pathway. This architecture enables robust learning of both fine-grained spectral variations and global patterns associated with drought stress. In this study, drought stress detection performance is evaluated by implementing and comparing two state-of-the-art models from the literature for tomato Vis–NIR spectral analysis^17,23^, along with three additional baseline AI models (CNN^21^, ResNet^25^, RF^22^) to ensure a comprehensive and fair comparative assessment. This experimental setup enables systematic benchmarking of the proposed method against both published approaches and standard machine learning and deep learning models.

In addition to evaluating Drought-Spec-Net using publicly available spectral data, we conducted an initial experiment using greenhouse RGB images collected at West Virginia State University. Although the original spectral study presented representative photographs of drought progression, the available dataset consisted of numerical Vis–NIR measurements and drought-stress labels without paired RGB images for individual spectral samples. To enable plant-level image analysis, we collected daily RGB images of drought-treated and well-watered control tomato plants. For this preliminary analysis, 44 plant images were selected for ilastik-based segmentation^26^ and quantification of visible canopy stress. This analysis produced an RGB-derived visible canopy stress index as a continuous image-based indicator of visible plant condition.

Beyond binary stress classification, we introduce a literature-informed potential yield-impact mapping that converts predicted drought probabilities into scenario-based impact indicators using a maximum impact level derived from previously reported tomato yield reductions. Because the spectral dataset does not contain measured yield outcomes, this mapping is intended to support exploratory interpretation and should not be considered a validated prediction of actual yield loss.

Finally, a PLLaMA-7B-Instruct-based agronomic reporting module was evaluated for converting drought probabilities, potential yield-impact indicators, and categorical stress levels into human-readable summaries. Together with the RGB-based quantification of visible canopy stress, these components establish an integrated framework combining spectral sensing, deep learning, potential yield-loss estimation, visible plant phenotyping, and automated reporting for practical decision support in agriculture. Fig. 1 illustrates the core spectral component of the framework, including Vis–NIR input, Drought-Spec-Net-based drought detection, potential yield-loss estimation, and LLM-based agronomic reporting.

**Figure 1.**
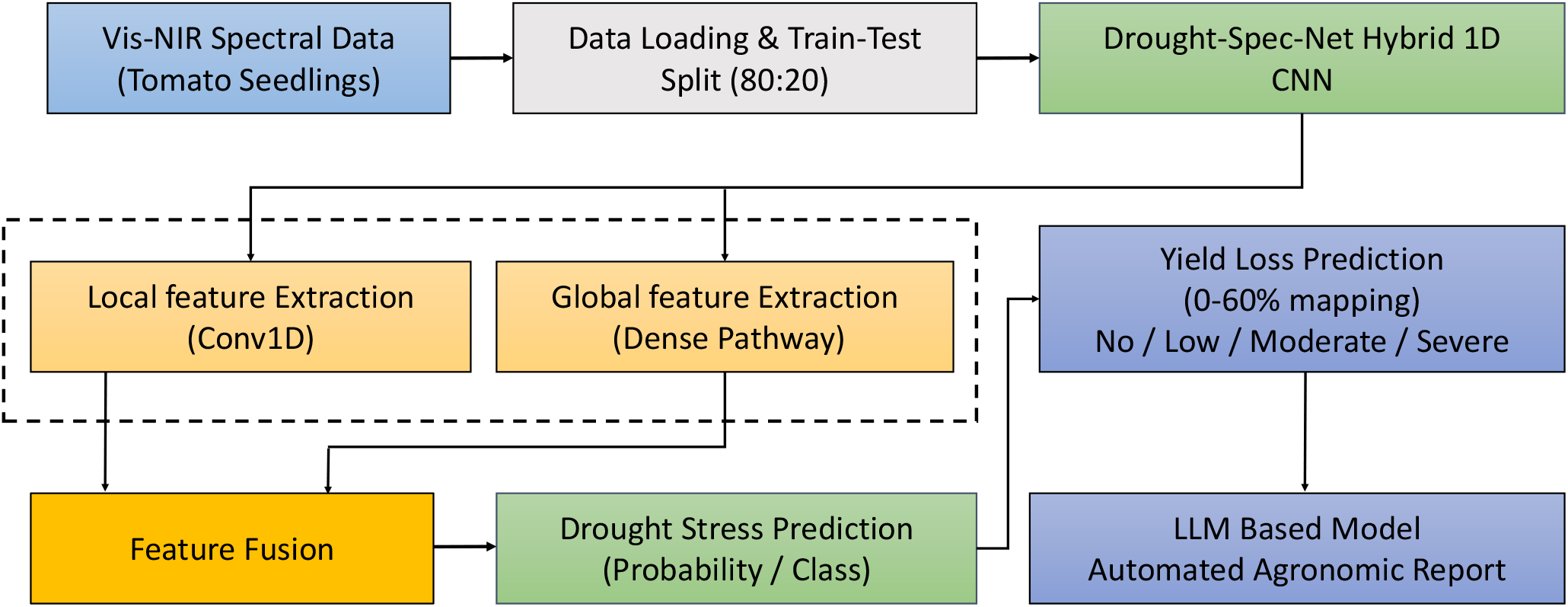
Overall framework of Drought-Spec-Net showing Vis–NIR spectral input, hybrid 1D CNN-based drought stress detection, potential yield-impact mapping, and LLM-based automated agronomic reporting for decision support in precision agriculture.

### Datasets and Experimental Design

This study comprises two complementary but independent experiments: (1) spectral drought-stress classification and literature-informed potential yield-impact mapping using a public Vis–NIR dataset and (2) visible canopy stress assessment using an independently collected WVSU greenhouse RGB image dataset. The two datasets were not paired, and the RGB images were not used as inputs to Drought-Spec-Net or to validate its spectral predictions.

#### Vis–NIR Spectral Dataset and Preparation

##### Vis–NIR Spectral Dataset

This study utilizes a publicly available visible–near-infrared (Vis–NIR) spectroscopic dataset of tomato seedlings for early drought stress detection. The dataset captures physiological responses of plants under well-irrigated and water-deficit conditions through high-resolution spectral measurements. It comprises 378 spectral samples, including 246 samples from well-irrigated (normal) plants and 132 samples from drought-stressed plants. The dataset was originally reported by Tu et al.^23^ and Kuo et al.^17^.

Each observation represents the averaged spectral signature of three fully expanded leaves from individual seedlings. Spectral data were acquired using an MS-720 portable spectroradiometer (EKO Instruments, San Jose, CA, USA), covering a wavelength range of 348–1052 nm. Each spectrum is discretized into 211 spectral bands at 3 nm intervals, forming a one-dimensional feature vector for each sample.

Each sample is annotated with an early-stage drought stress label, where 0 denotes normal conditions and 1 indicates drought stress. These spectral measurements capture variations in plant water content, pigment composition, and cellular structure, enabling the detection of drought stress prior to visible morphological symptoms.

Figure 2 illustrates the mean spectral reflectance profiles of normal and drought-stressed tomato seedlings across the Vis–NIR wavelength range (348–1052 nm). A clear distinction between the two classes can be observed across multiple spectral regions, indicating that drought stress induces measurable changes in plant reflectance characteristics. These variations demonstrate that the spectral signatures contain discriminative information for differentiating between healthy and stressed plants, supporting the suitability of the dataset for data-driven drought stress detection.

**Figure 2.**
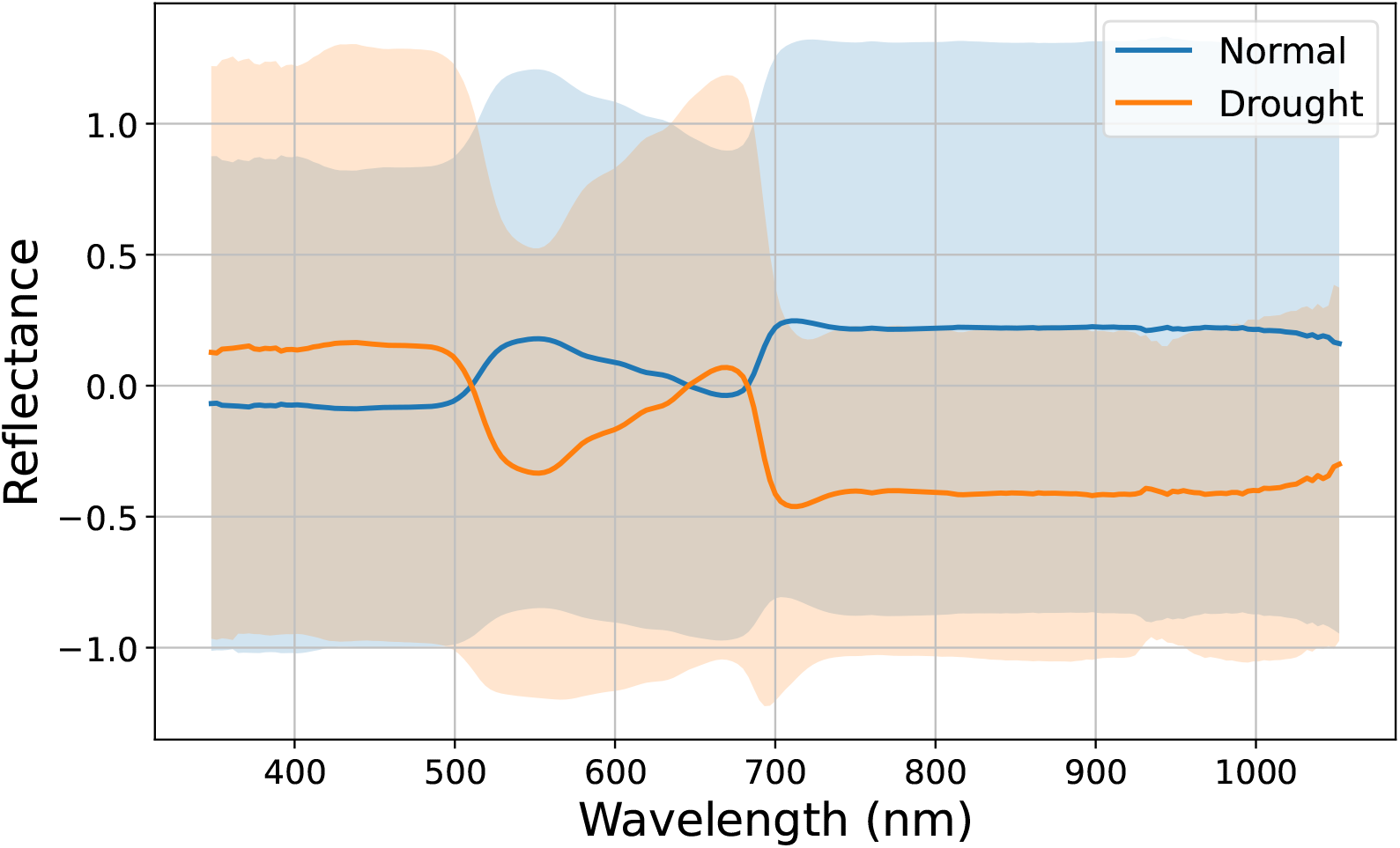
Mean spectral reflectance profiles of healthy and drought-stressed tomato seedlings across the Vis–NIR spectral range (348–1052 nm).

The imbalance ratio (IR) is defined as the ratio between the number of majority class samples and minority class samples, where IR = 2 corresponds to the original dataset with natural class distribution (246 normal and 132 drought samples, as described above). To further analyze model robustness, a balanced dataset (IR = 1) is constructed through controlled resampling of the original dataset, resulting in 189 samples for each class. In addition, datasets with IR = 10 and IR = 50 are generated from the original dataset using controlled random sampling, yielding 340 normal and 38 drought samples for IR = 10, and 370 normal and 8 drought samples for IR = 50. This setup enables evaluation under progressively varying class imbalance conditions under a unified experimental protocol^23^.

##### Spectral Data Preparation

The dataset consists of one-dimensional visible–near-infrared (Vis–NIR) spectral signals, where each sample is represented by 211 wavelength features. These spectral samples are directly used as input for model development.

The dataset was divided into training and testing sets using an 80:20 split. From the training subset, 20% was further allocated for validation during model training, resulting in an effective partitioning of 64% training, 16% validation, and 20% testing data. The validation set was used to monitor training performance and support model optimization, while the test set was kept completely unseen until final evaluation to ensure unbiased assessment and prevent data leakage. Each sample was reshaped into a format compatible with the one-dimensional hybrid convolutional neural network used in this study.

#### WVSU Greenhouse Experiment and RGB Dataset

##### Plant Materials and Experimental Design

Tomato (*Solanum lycopersicum*) introgression lines (ILs), derived from a cross between the cultivated tomato cv. M82 and the wild relative S. pennellii, were evaluated for drought responses under greenhouse conditions at West Virginia State University from June to August 2026. Uniform, healthy five-week-old seedlings were transplanted individually into 3-gallon pots containing Pro-Mix growing medium and placed on the PlantArray phenotyping platform (Plant-DiTech, Israel). The experiment was arranged in a randomized complete block design (RCBD) with three biological replicates per treatment. Each pot was connected to an individual irrigation unit, and the growing medium was initially saturated and allowed to drain freely to establish uniform, well-watered conditions. Following transplantation, all plants were maintained under well-watered conditions for 10 days to allow plant establishment and acclimation to the PlantArray system. During the 10-day well-watered acclimation period, pots were irrigated at night using 3-min irrigation events at 15-min intervals for 4 h, with dripper assemblies positioned within the growing medium to ensure uniform wetting and maintain non-limiting water conditions. After the 10-day acclimation period, plants were assigned to well-watered control and drought treatments. Well-watered plants continued to receive the same irrigation regime, whereas irrigation was completely withheld from drought-treated plants to induce progressive soil drying.

##### RGB Image Dataset

RGB images of individual drought-treated and well-watered control tomato plants were acquired daily throughout the 20-day drought-treatment period at the West Virginia State University greenhouse to document plant development and the progression of visible drought symptoms. The greenhouse setup and plant arrangement are shown in Fig. 3a. For this preliminary analysis, 44 images, each representing a different individual plant, were randomly selected from the daily image collection across both treatment groups for pixel-level classification and visible canopy-stress quantification. A representative plant image and its corresponding three-class mask are shown in Fig. 3b,c.

**Figure 3.**
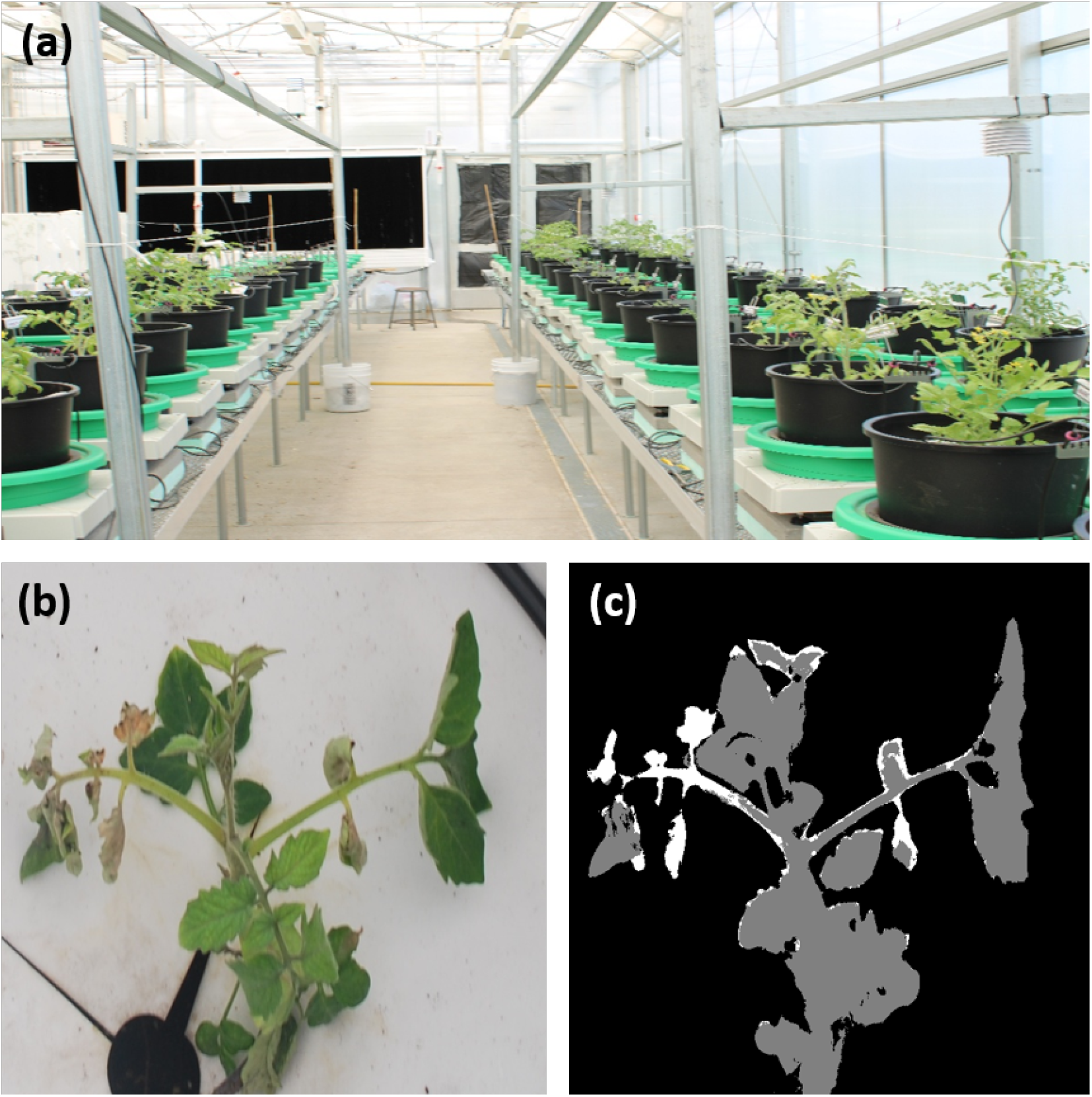
WVSU greenhouse RGB dataset and canopy segmentation. (a) Greenhouse experimental setup containing drought-treated and well-watered control tomato plants. (b) Representative individual-plant RGB image selected from the daily image collection. (c) Corresponding three-class mask generated using ilastik, where black represents background, gray represents healthy-green canopy tissue, and white represents visibly drought-stressed canopy tissue.

### Methodology

#### Proposed Drought-Spec-Net Architecture

The proposed Drought-Spec-Net is a hybrid one-dimensional convolutional neural network designed for early drought stress detection using Vis–NIR spectral data. The model takes a one-dimensional input vector of length 211, representing spectral reflectance values across the visible–near-infrared wavelength range.

The architecture is composed of two complementary feature extraction branches (See Figure 4). The first branch is a convolutional pathway that captures local spectral patterns through stacked 1D convolutional layers followed by ReLU activation and max-pooling operations. This branch enables the model to learn fine-grained spectral variations associated with drought-induced physiological changes.

**Figure 4.**
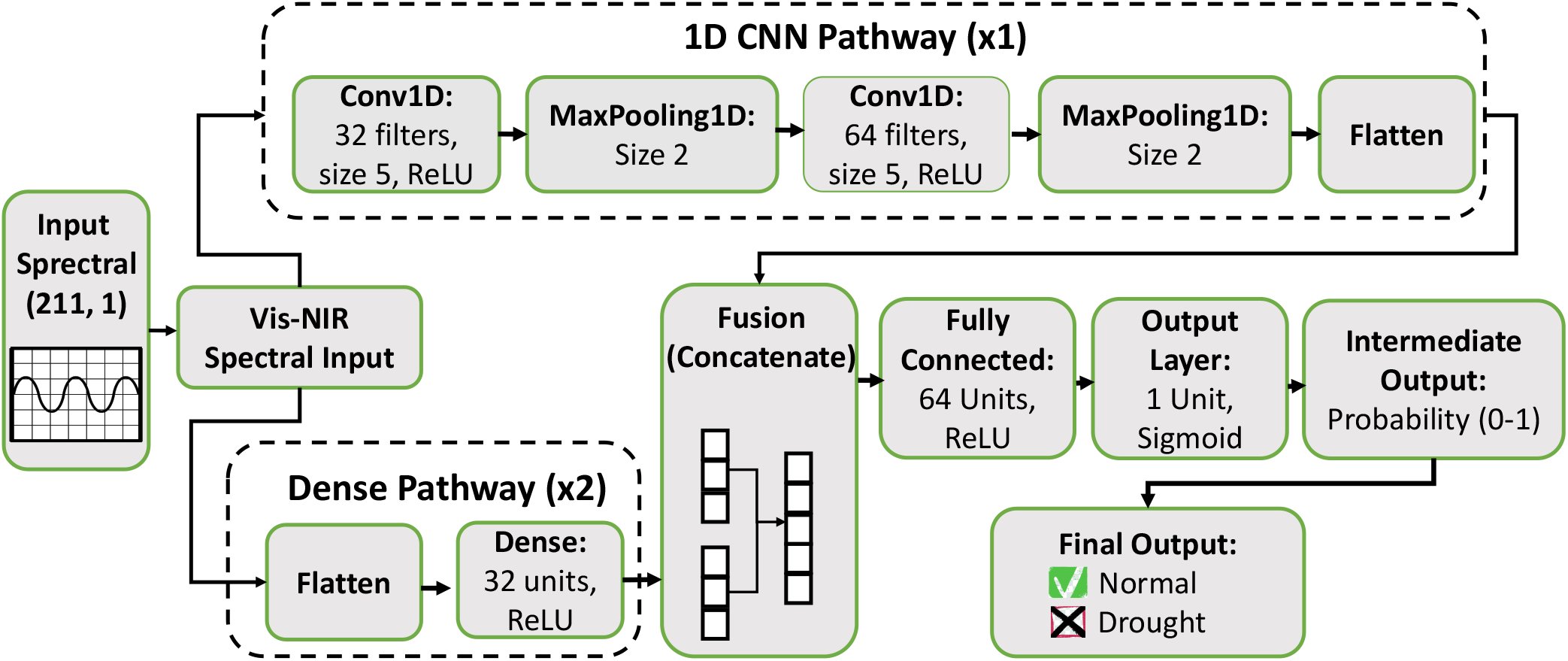
Architecture of the proposed DroughtSpecNet model. The input Vis–NIR spectral signal is processed through a hybrid framework consisting of a 1D convolutional pathway for local feature extraction and a parallel dense pathway for global feature learning. The outputs of both branches are concatenated and passed through a fully connected layer, followed by a sigmoid activation function to produce a probability score. A threshold-based decision rule (*p≥* 0.5) is applied to perform binary classification of drought stress.

The second branch is a parallel dense pathway that operates directly on the flattened spectral input. This branch preserves global spectral information and helps capture overall trends in the reflectance signal that may not be fully captured by convolutional filters.

The outputs from both branches are concatenated to form a fused feature representation, combining local discriminative patterns with global contextual information. The fused features are then passed through a fully connected dense layer to learn higher-level representations.

Finally, the model uses a sigmoid activation function in the output layer to perform binary classification, distinguishing between normal and drought-stressed samples. This hybrid design improves feature representation capability by integrating convolution-based local feature extraction with dense global feature learning, making it well-suited for complex spectral classification tasks.

#### Comparative Models

To evaluate the performance of the proposed Drought-Spec-Net, five comparative models were selected, including two state-of-the-art (SOTA) methods and three baseline models. All deep learning models were trained under identical experimental settings, including 200 epochs and a batch size of 16, while using the same training, validation, and testing splits to ensure a fair, consistent, and unbiased comparison across all methods.

The selected models represent both recent deep learning advancements in spectral-based plant stress analysis and a classical machine learning approach. This combination allows a comprehensive evaluation of performance across different levels of model complexity, ranging from conventional feature learning to advanced deep neural architectures with attention mechanisms

- One Dimension Spectrogram Power Net (1D-SP-Net^23^) The 1D-SP-Net model was included as a state-of-the-art baseline for comparison. Although the authors provided a GitHub repository, it did not include a complete training pipeline suitable for direct reproduction. Therefore, the model was reimplemented based on the detailed architectural description provided in the original publication. The architecture integrates residual learning and global context (GC) modules to capture both local and global spectral dependencies. It also incorporates channel-wise attention, batch normalization, dropout, and swish activation functions to improve feature representation and generalization. The model was evaluated under the same experimental settings as the proposed approach.
- 1D CNN with an embedded residual global context Block (1D-ResGC-Net^17^) A 1D-Res-Net with attention mechanism was used as a state-of-the-art baseline for comparison. The model is based on a residual learning framework enhanced with channel-wise attention to improve feature representation from spectral signals. The architecture consists of stacked 1D convolutional layers with residual connections, batch normalization, and dropout for regularization. An attention module is applied after each residual block using global average pooling followed by learnable channel re-weighting. Swish activation functions are used throughout the network to improve non-linear feature learning. Since the publicly available resources did not provide a complete and directly executable implementation, the model was reimplemented based on the architectural description from the original work and associated materials. The model was trained under the same experimental setup as the proposed method.
- 1D Convolutional Neural Network (CNN) A standard 1D-CNN was implemented as a baseline deep learning model. It consists of stacked convolutional layers with max-pooling operations for feature extraction and dimensionality reduction. The extracted features are passed through a fully connected layer followed by a sigmoid output layer for binary classification of drought stress.
- 1D Residual Network (ResNet) A conventional 1D-Res-Net model was used to evaluate the impact of residual learning without additional enhancements such as attention mechanisms. The architecture consists of convolutional layers with skip connections that improve gradient flow and enable stable training of deeper networks.
- Random Forest (RF)

A Random Forest classifier with 200 decision trees was used as a classical machine learning baseline. The model constructs multiple decision trees using bootstrapped subsets of the training data and random feature selection. Final predictions are obtained through majority voting across all trees. This model provides a non-deep learning benchmark for evaluating the effectiveness of spectral feature learning approaches.

### Training Strategy

All deep learning models, including the proposed Drought-Spec-Net, 1D-CNN, 1D-Res-Net, 1D-ResGC-Net, and the 1D-SP-Net, were trained under a consistent experimental protocol to ensure a fair and unbiased comparison. Each model was trained for 200 epochs with a batch size of 16. All models were trained and evaluated using the same training, validation, and test partitions described in the Spectral Data Preparation subsection. A fixed random seed (`random_state = 42`) was used to ensure reproducibility and consistent data partitions across all models.

All deep learning models were optimized using the Adam optimizer with binary cross-entropy loss, which is suitable for binary classification tasks such as drought stress detection. Model weights were updated iteratively using backpropagation to minimize the loss function over the training epochs. To improve training stability and generalization, techniques such as batch normalization and dropout were incorporated within the network architectures. These components help reduce overfitting and improve robustness when learning from high-dimensional spectral data.

For the Random Forest model, the same training and testing splits were used to ensure consistency. The model was trained using an ensemble of 200 decision trees, and no deep learning-specific optimization strategies were applied.

Overall, this unified training strategy ensures that performance differences among models are primarily due to architectural design rather than differences in training conditions.

### Literature-Informed Potential Yield-Loss Mapping

Drought stress is defined as an altered physiological condition that disrupts the homeostatic balance of plants, affecting their metabolic processes and growth^27,28^. Among abiotic stresses, drought is considered the most severe and pervasive factor limiting agricultural productivity. It arises due to insufficient water availability caused by irregular precipitation, high evapotranspiration, and limited soil moisture retention, which collectively prevent plants from achieving their full genetic yield potential^29^. Water deficit significantly impacts key physiological processes, particularly photosynthesis, leading to cellular dehydration, reduced biomass accumulation, and ultimately decreased crop yield. Previous studies indicate that drought stress can cause substantial yield losses across crops, especially in tomato, with reductions of up to 55% overall and as high as 70% in highly susceptible genotypes under severe conditions^8,13,30,31^.

In this study, the drought probabilities generated by Drought-Spec-Net from Vis–NIR spectral data were used to calculate a potential yield-impact indicator. Because the dataset contains only binary labels for normal and drought-stressed samples and does not include measured plant-level yield, the model probabilities represent confidence in drought classification rather than directly measured drought severity. Therefore, a literature-informed mapping function was applied to probabilities above the classification threshold to generate exploratory, scenario-based estimates of potential yield impact.

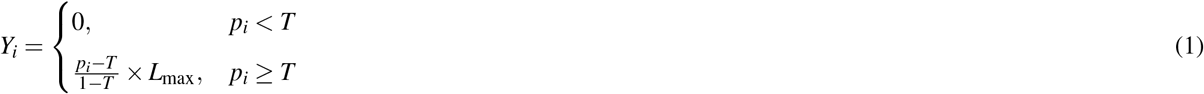

where *Y*_*i*_ represents the potential yield-impact indicator for sample *i* (%), *p*_*i*_ is the predicted drought probability for sample *i, T* = 0.5 is the threshold separating healthy and stressed conditions, and *L*_max_ is the assumed maximum potential impact under severe drought conditions.

Since no direct ground-truth yield measurements are available, this formulation provides an approximate yet interpretable estimation of yield loss. Based on prior agronomic studies, drought stress in tomato can result in yield reductions of approximately 55%, with losses typically reaching up to 70% under severe drought conditions. To ensure a realistic and conservative estimate within this reported range, the maximum potential yield-impact level was set to *L*_max_ = 60%.

For interpretability, the estimated yield loss values are further categorized into four levels of severity: no loss (0%), low loss (0–25% of *L*_max_), moderate loss (25–75% of *L*_max_), and severe loss (*>* 75% of *L*_max_). This categorization facilitates intuitive interpretation of drought severity and its potential impact on crop productivity, providing a practical linkage between model predictions and agronomic decision-making.

### RGB Pixel Classification and Visible Canopy Stress Quantification

The 44 selected RGB images were processed using the pixel-classification workflow in ilastik. Pixels were assigned to three classes: background, healthy-green canopy tissue, and visibly drought-stressed canopy tissue. The resulting masks were exported as grayscale PNG images, with pixel values of 0, 128, and 255 representing background, healthy-green tissue, and visibly stressed tissue, respectively. Each mask was checked for unexpected pixel values and the presence of canopy pixels before analysis.

For plant image *i*, the total canopy pixel count was calculated as

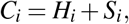

where *H*_*i*_ and *S*_*i*_ represent the numbers of healthy-green and visibly stressed canopy pixels, respectively. Background pixels were excluded from all canopy-level calculations. The healthy fraction (*HF*) and RGB-derived drought severity index (*DSI*) were calculated as

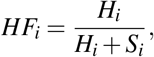

and

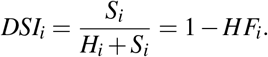

The DSI ranges from 0 to 1, where values closer to 0 indicate predominantly healthy-green canopy tissue and values closer to 1 indicate a greater proportion of visibly stressed canopy tissue. Each plant image, rather than each pixel, was treated as one observational unit. Plant-level HF and DSI values were summarized using the mean, standard deviation, median, interquartile range, minimum, and maximum. A stacked canopy-composition plot and a histogram were generated to visualize plant-level healthy-green and visibly stressed fractions and the overall DSI distribution, respectively. The RGB-derived DSI was interpreted only as an indicator of visible canopy stress and was not directly converted into yield loss because corresponding plant-level yield measurements were unavailable.

### LLM-Based Agronomic Reporting

To improve interpretability and increase usability of drought stress predictions, an LLM-based agronomic reporting module, named AgriLLaMA, was integrated into the proposed framework. While Drought-Spec-Net provides probabilistic drought stress predictions and estimated yield loss values, these outputs alone may not be sufficiently interpretable for practical agricultural decision-making. Therefore, an LLM layer was introduced to automatically transform numerical outputs into human-readable agronomic summaries and management recommendations.

In this study, PLLaMa-7B-Instruct^32^, a domain-specific large language model optimized for plant science applications, was further adapted and utilized to develop AgriLLaMA for automated agronomic report generation. PLLaMa-7B-Instruct is based on the LLaMA-2 architecture and was further pretrained on more than 1.5 million plant science academic articles followed by instruction tuning for agricultural question answering and text generation tasks^32^. The model is used locally using the Hugging Face Transformers framework and executed on NVIDIA RTX PRO 4500 GPUs. Two GPUs each having 32 GigaByte of GDDR7 VRAM are used for this purpose.

The reporting framework is designed to receive outputs from the drought detection and yield estimation pipeline, including predicted drought probability, estimated yield loss percentage, and categorical stress severity. These variables are then incorporated into prompting templates to generate agronomic interpretations and irrigation recommendations.

To analyze the impact of prompt engineering and hallucination control, three different prompting strategies were evaluated using 60 synthetic agronomic samples evenly balanced across Low, Moderate, and Severe drought stress categories. Each sample was evaluated using all three prompting strategies, resulting in a total of 180 generated outputs. The evaluated prompting configurations are summarized as follows:

- **Baseline Prompt:** The baseline setting directly provided raw numerical outputs to AgriLLaMA without structural guidance. This setting evaluates unconstrained generative behavior using only drought probability and estimated yield loss values. For example: “A tomato plant has drought probability = 0.82 and estimated yield loss = 38
- **Semi-Guided Prompt:** The semi-guided setting introduced predefined report sections including stress level, estimated yield loss, interpretation, and recommendation while still allowing flexible natural language generation. For example: “Generate an agronomic report using the following sections: Stress Level, Estimated Yield Loss, Interpretation, and Recommendation. Use only the provided crop, drought probability, and estimated yield loss values.”
- **Strictly Controlled Prompt:** The strictly controlled setting enforced explicit factual constraints and JSON output formatting to minimize hallucination and improve factual consistency. The model was instructed to preserve all numerical values exactly, avoid introducing unsupported information, and generate outputs using a predefined JSON schema containing stress level, risk interpretation, recommendation, irrigation strategy, confidence level, and justification fields. For example: “Return only valid JSON. Do not invent values. Preserve all numerical values exactly. Use only the provided inputs.”

### Evaluation Metrics

The performance of the proposed drought detection model was evaluated using standard classification metrics, including accuracy, precision, recall, and F1-score. These metrics provide complementary insights into model performance, particularly in handling class imbalance between normal and drought-stressed samples.

Accuracy measures the overall correctness of predictions, while precision quantifies the proportion of correctly predicted drought-stressed samples among all predicted stressed instances. Recall evaluates the model’s ability to correctly identify actual drought-stressed samples, and the F1-score provides a balanced combination of precision and recall, offering a balanced assessment of model performance.

In addition, a confusion matrix was used to provide a detailed breakdown of prediction outcomes, including true positives, true negatives, false positives, and false negatives, enabling a comprehensive understanding of classification behavior. To monitor training dynamics and assess model generalization, epoch-wise training and validation accuracy and loss were tracked throughout the training process. These curves provide insight into model convergence, stability, and potential overfitting or underfitting behavior.

For the AgriLLaMA-based agronomic reporting module, we evaluated using factual consistency, numerical preservation, hallucination behavior for different prompting strategies. Category consistency was used to evaluate whether the generated report preserved the provided drought severity category information. Numerical consistency measured whether drought probability and estimated yield loss values were preserved without modification or omission.

Hallucination occurrence was evaluated by identifying unsupported or fabricated content introduced by the language model that was not present in the input prompt. For the strictly controlled prompting setting, structural compliance was additionally evaluated based on adherence to the predefined JSON output schema.

## Results and Discussion

### Performance of Drought Detection Models

The results reported in this section are obtained using the original dataset. The performance of the proposed model was evaluated using multiple quantitative metrics, including accuracy, precision, recall, and F1-score, along with training dynamics and ROC analysis.

The learning behavior of the model is illustrated in Fig. 5, which presents the training and validation accuracy and loss curves over epochs. The model demonstrates rapid convergence during the initial training phase, with training accuracy increasing steadily and reaching approximately 97–98%. The validation accuracy follows a similar trend, stabilizing around 88–91%. The loss curves further confirm effective learning, where training loss decreases consistently across epochs. In contrast, the validation loss shows moderate fluctuations but still maintaining good generalization performance. Overall, the model achieves stable convergence without significant degradation in validation performance.

**Figure 5.**
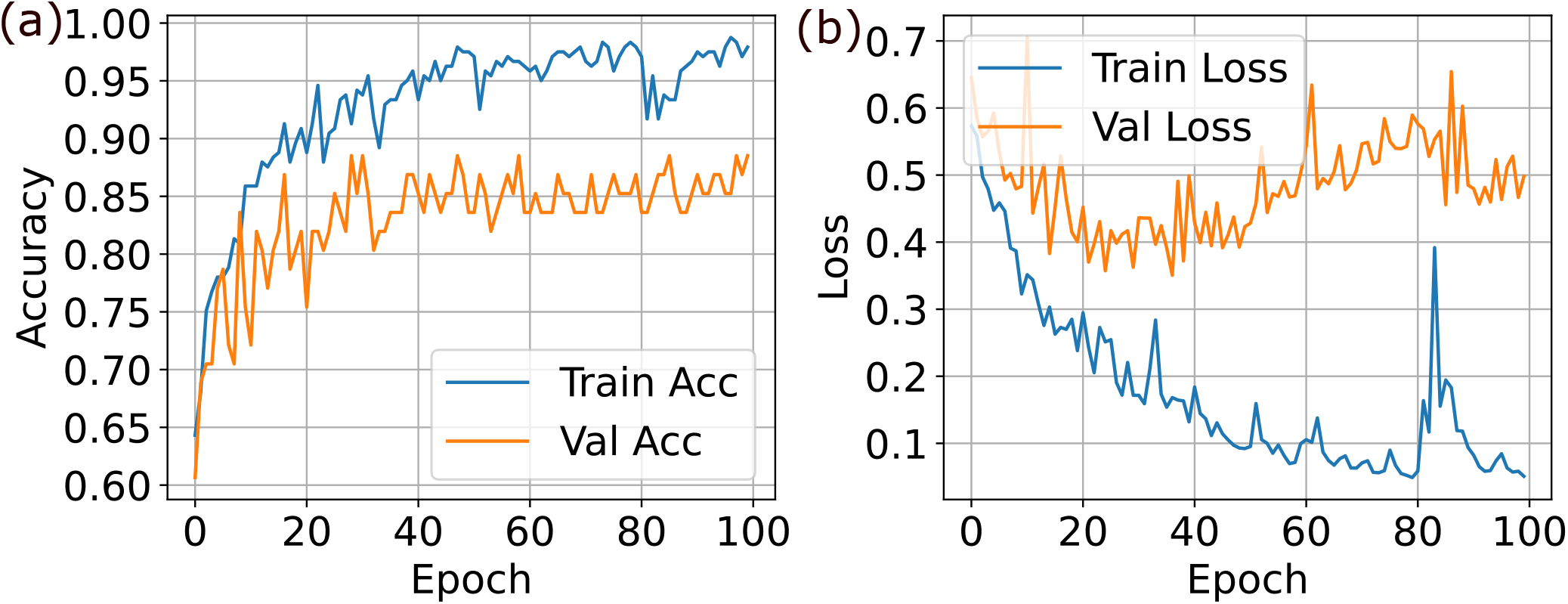
Training and validation performance of the proposed model: (a) training and validation accuracy, and (b) training and validation loss.

On the test dataset, the proposed model achieves an accuracy of 97.37%, demonstrating its effectiveness in drought stress classification. The model attains a precision of 95.24%, recall of 98.25%, and an F1-score of 96.61%, indicating a strong balance between precision and recall. The high recall value is particularly important, as it reflects the model’s ability to correctly identify drought stress conditions with minimal false negatives.

A detailed classification report further highlights class-wise performance. For the *Normal* class, the model achieves a precision of 100.00% and recall of 96.00%, resulting in an F1-score of 98.00%. For the *Drought* class, the precision is 90.00% and recall reaches 100.00%, yielding an F1-score of 95.00%. These results indicate that the model is highly effective in detecting drought conditions while maintaining strong overall classification performance.

The discriminative capability of the model is further validated using the Receiver Operating Characteristic (ROC) curve, as shown in Fig. 6 (a). The model achieves an Area Under the Curve (AUC) of 99.63%, indicating excellent separability between normal and drought stress conditions.

**Figure 6.**
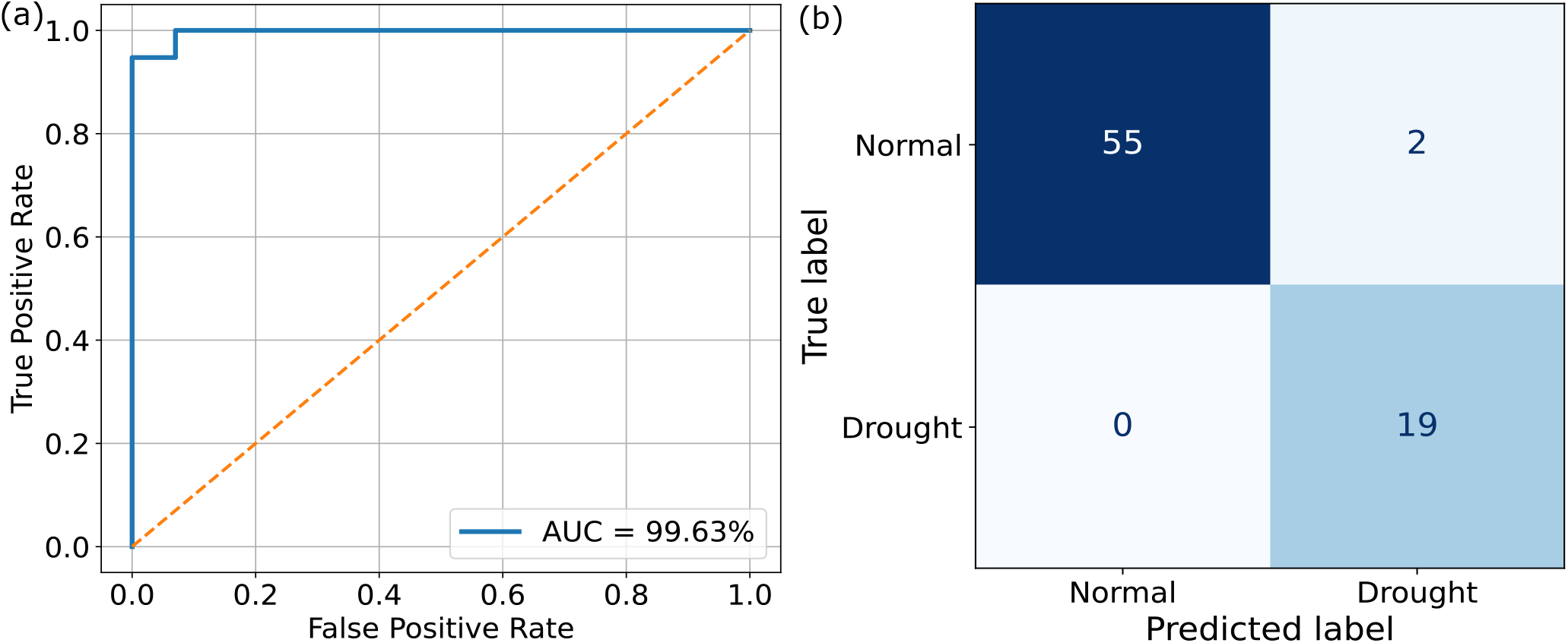
Performance evaluation of the proposed Drought-Spec-Net on the test dataset: (a) ROC curve with an AUC of 99.63%, and (b) Confusion matrix illustrating correct and incorrect classifications for normal and drought-stressed samples..

These results confirm that the proposed model provides robust and reliable performance for drought stress detection under the given experimental setup.

### Spectral Discrimination between Normal and Drought-Stressed Samples

To further evaluate the effectiveness of the proposed model in distinguishing between normal and drought stress conditions, a confusion matrix analysis was conducted, as shown in Fig. 6 (b).

The confusion matrix reveals that out of 57 normal samples, 55 were correctly classified, while 2 samples were misclassified as drought, resulting in a true negative rate of 96.49%. For the drought class, all 19 samples were correctly identified, achieving a true positive rate (recall) of 100.00%. Notably, the model produces zero false negatives, which is highly desirable in drought detection applications where missing stress conditions can have significant consequences.

The precision for the drought class is 90.48%, indicating that a small number of normal samples are incorrectly predicted as drought. This slight tendency toward false positives suggests that the model is more sensitive to detecting drought conditions, prioritizing recall over precision. Such behavior is often preferred in agricultural monitoring systems, where early detection of stress is critical.

Overall, the confusion matrix demonstrates that the proposed model achieves strong class-wise discrimination, with high accuracy in both normal and drought categories. The results confirm that the model is particularly effective in identifying drought stress, making it suitable for reliable deployment in precision agriculture applications.

### Comparative Analysis with State-of-the-Art Methods

To validate the effectiveness of the proposed model, a comparative analysis was conducted against several baseline and state-of-the-art models, including 1D-SP-Net, 1D-ResGC-Net, standard 1D-CNN, 1D-Res-Net, and a classical Random Forest (RF) model. All models were evaluated under identical experimental conditions.

As shown in Table 1, the proposed model outperforms all comparative methods across all evaluation metrics. Specifically, it achieves the highest accuracy of 97.37%, significantly improving upon the baseline 1D-CNN (93.42%) and other deep learning architectures such as 1D-Res-Net and 1D-ResGC-Net.

**Table 1.** Comparative performance of different models for drought stress classification.

| Model | Accuracy (%) | Precision (%) | Recall (%) | F1-score (%) |
| --- | --- | --- | --- | --- |
| 1D-SP-Net | 90.79 | 75.00 | 94.74 | 83.72 |
| 1D-ResGC-Net | 89.47 | 78.95 | 78.95 | 78.95 |
| 1D-CNN | 93.42 | 89.98 | 93.86 | 91.65 |
| 1D-Res-Net | 89.47 | 73.91 | 89.47 | 80.95 |
| RF | 77.63 | 71.31 | 74.56 | 72.43 |
| **Proposed Drought-Spec-Net** | **97.37** | **95.24** | **98.25** | **96.61** |

In terms of precision, the proposed model reaches 95.24%, indicating fewer false positive predictions compared to other models, particularly 1D-SP-Net (75.00%) and RF (71.31%). The recall of 98.25% is the highest among all models, demonstrating superior capability in correctly identifying drought stress conditions.

The F1-score of 96.61% further confirms the robustness of the proposed model by maintaining a strong balance between precision and recall. In contrast, other models show either lower precision or recall, leading to reduced overall performance.

Traditional machine learning approaches such as RF exhibit significantly lower performance, highlighting the advantage of deep learning-based feature extraction. Even among deep learning models, the proposed architecture consistently delivers superior results, demonstrating its effectiveness for drought stress classification.

Overall, the results confirm that the proposed model provides a substantial improvement over existing methods and is highly suitable for accurate and reliable drought stress detection.

### Robustness Analysis under Varying Imbalance Ratios

While the previous analyses are conducted on the original dataset, additional experiments are performed to assess the robustness of the proposed model under varying class imbalance conditions.

In this study, three additional dataset variants are constructed corresponding to imbalance ratios of 1, 10, and 50, representing balanced, moderate, and severe class imbalance scenarios, respectively. The balanced dataset (IR = 1) is obtained through controlled resampling of the original dataset, while the more imbalanced settings (IR = 10 and IR = 50) are generated from the original dataset via controlled random sampling of majority and minority class samples to simulate progressively skewed class distributions, following the experimental protocol described in^23^.

All baseline, state-of-the-art, and proposed models are evaluated under identical dataset splits and experimental protocols across all imbalance conditions to ensure a fair comparison. This setup enables a systematic assessment of model stability and generalization performance under diverse class distribution scenarios.

The detailed comparative results across all imbalance settings are summarized in Table 2. The performance trends clearly demonstrate how different models behave under varying class distribution complexities, highlighting both stability and degradation patterns across increasing imbalance ratios.

**Table 2.** Performance comparison of all models across different imbalance ratios (IR)

| Dataset (IR) | Model | Accuracy (%) | Precision (%) | Recall (%) | F1-score (%) |
| --- | --- | --- | --- | --- | --- |
| IR = 1 (Balanced) | 1D-SP-Net | 90.79 | 95.12 | 88.64 | 91.76 |
|  | 1D-ResGC-Net | 88.16 | 88.89 | 90.91 | 89.89 |
|  | 1D-CNN | 93.42 | 93.07 | 93.89 | 93.33 |
|  | 1D-Res-Net | 89.47 | 97.37 | 84.09 | 90.24 |
|  | RF | 84.21 | 83.75 | 84.23 | 83.93 |
|  | Proposed Drought-Spec-Net | **93.42** | **93.13** | **93.47** | **93.28** |
| IR = 2 (Original) | 1D-SP-Net | 90.79 | 75.00 | 94.74 | 83.72 |
|  | 1D-ResGC-Net | 89.47 | 78.95 | 78.95 | 78.95 |
|  | 1D-CNN | 93.42 | 89.98 | 93.86 | 91.65 |
|  | 1D-Res-Net | 89.47 | 73.91 | 89.47 | 80.95 |
|  | RF | 77.63 | 71.31 | 74.56 | 72.43 |
|  | Proposed Drought-Spec-Net | **97.37** | **95.24** | **98.25** | **96.61** |
| IR = 10 (Moderate) | 1D-SP-Net | 94.74 | 87.50 | 70.00 | 77.78 |
|  | 1D-ResGC-Net | 89.47 | 55.56 | 100.00 | 71.43 |
|  | 1D-CNN | 94.74 | 91.54 | 84.24 | 87.40 |
|  | 1D-Res-Net | 96.05 | 100.00 | 70.00 | 82.35 |
|  | RF | 89.47 | 94.59 | 60.00 | 63.81 |
|  | Proposed Drought-Spec-Net | **96.05** | **97.83** | **85.00** | **90.07** |
| IR = 50 (Severe) | 1D-SP-Net | 96.05 | 0.00 | 0.00 | 0.00 |
|  | 1D-ResGC-Net | 23.68 | 4.92 | 100.00 | 9.38 |
|  | 1D-CNN | 96.05 | 48.03 | 50.00 | 48.99 |
|  | 1D-Res-Net | 96.05 | 0.00 | 0.00 | 0.00 |
|  | RF | 96.05 | 48.03 | 50.00 | 48.99 |
|  | Proposed Drought-Spec-Net | **96.05** | **48.03** | **50.00** | **48.99** |

From Table 2, it can be observed that the performance under the balanced setting (IR = 1) and the original dataset (IR = 2) remains relatively close across most models. In some cases, the proposed model achieves slightly higher performance on the original dataset, particularly in terms of recall and F1-score, indicating that the model is well-adapted to the natural data distribution. Overall, both IR = 1 and IR = 2 settings show comparable behavior, suggesting that the proposed framework is stable even when the class distribution is slightly adjusted.

For moderate imbalance (IR = 10), a noticeable variation in model behavior is observed. While deep learning-based models such as 1D-CNN and the proposed Drought-Spec-Net maintain relatively strong performance, traditional architectures and baseline models begin to show reduced recall, indicating difficulty in capturing minority class patterns under skewed distributions. In particular, the Random Forest model exhibits a significant drop in recall, highlighting its sensitivity to class imbalance.

Under severe imbalance conditions (IR = 50), a substantial degradation in performance is observed across most baseline models. Models such as 1D-SP-Net and 1D-Res-Net show collapse in precision and F1-score, indicating failure to reliably identify the minority class. Although some models maintain relatively stable accuracy, this is largely due to bias toward the majority class, which does not reflect meaningful classification performance. In contrast, the proposed model retains comparatively more stable behavior, although a reduction in F1-score is still observed, reflecting the extreme difficulty of learning under highly imbalanced conditions.

Overall, these results confirm that while all models perform reasonably well under mild imbalance, their robustness decreases as the imbalance ratio increases. The proposed Drought-Spec-Net demonstrates better stability compared to baseline and state-of-the-art methods, particularly in maintaining balanced precision and recall under challenging data distributions.

### Potential Yield-Loss Mapping Results

The yield loss estimation results are obtained using the original dataset with its natural class distribution. The estimated yield loss distribution derived from the Drought-Spec-Net predictions demonstrates a wide variability in drought-induced productivity reduction across the dataset. Overall, the predicted yield loss values range from 0% to 60%, with a maximum loss of 60% and a minimum loss of 0%, indicating that both fully healthy and severely stressed conditions are captured by the model. The average predicted yield loss is 12.97%, suggesting that most samples fall within low to moderate stress conditions.

A detailed statistical breakdown shows a highly skewed distribution toward minimal yield impact. The majority of samples, 55 out of 76 (72.37%), fall under the No Loss (0%) category, indicating healthy or near-healthy plants with negligible predicted productivity reduction. A small number of samples, 1 (1.32%), fall under the Low loss category, showing minimal stress influence.

In contrast, Severe yield loss accounts for 12 samples (15.79%), representing plants experiencing high drought stress with substantial predicted productivity reduction. Additionally, 8 samples (10.53%) are classified under Moderate loss, indicating intermediate stress conditions with noticeable but not extreme yield impact.

The yield loss distribution is further illustrated in the histogram of predicted values (Fig. 7 (a)), which shows a strong concentration of predictions near 0%, with a secondary cluster approaching the upper bound near 60%. This pattern reflects the model’s tendency to classify most samples as either healthy or strongly stressed, with fewer intermediate cases.

**Figure 7.**
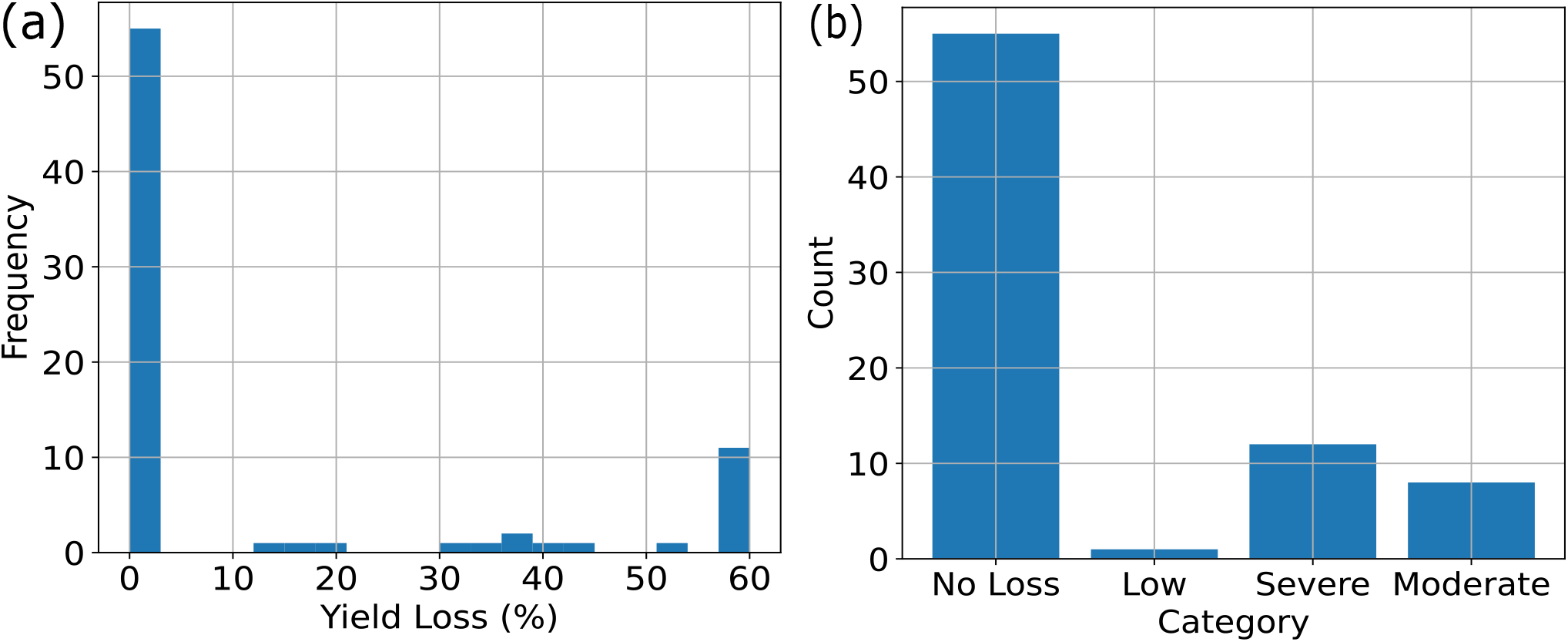
Potential yield-impact mapping derived from Drought-Spec-Net probabilities. (a) Distribution of predicted yield loss values showing the frequency of samples across the full loss range. (b) Corresponding categorical distribution of yield loss levels including No Loss, Low, Moderate, and Severe classes.

The categorical breakdown of yield loss levels is shown in (Fig. 7 (b)), which confirms that the dataset is dominated by low-stress conditions while a smaller subset exhibits moderate to severe drought impact.

Overall, these results demonstrate that the proposed framework not only performs accurate drought stress classification but also provides meaningful agronomic quantification of potential yield losses, enabling practical interpretation for precision irrigation and crop management strategies.

### AgriLLaMA-Based Agronomic Reporting Results

A total of 60 synthetic agronomic samples were evaluated using three different prompting strategies, resulting in 180 generated outputs. Experimental observations revealed substantial behavioral differences across prompting strategies in terms of factual consistency, numerical preservation, hallucination occurrence, and structural reliability.

*Baseline prompting* frequently produced verbose and inconsistent outputs containing fabricated contextual information unrelated to the provided drought prediction inputs. Several generated responses included unsupported environmental reasoning, fabricated recipients, unrelated narrative structures, and agronomic assumptions not present in the input prompt. Representative hallucination examples included generated phrases such as “Dear [Recipient’s Name]”, unsupported environmental reasoning, and fabricated explanatory content unrelated to drought probability or estimated yield loss. *Semi-guided prompting* reduced verbosity and partially improved topical relevance; however, the generated outputs still exhibited unsupported agronomic interpretations and repetitive stress descriptions. In several cases, numerical drought probability and yield loss values were omitted entirely from the generated reports despite being provided in the prompt. *Strictly controlled* prompting significantly improved structural consistency and category preservation behavior compared to baseline and semi-guided prompting. The generated outputs demonstrated substantially improved adherence to provided drought severity categories. However, malformed JSON structures, instruction leakage, schema drift, and partially fabricated reasoning were still observed in multiple outputs despite explicit formatting constraints.

Table 3 summarizes the quantitative evaluation results across all prompting strategies. Category consistency measured whether the generated output preserved the provided drought severity category. Probability consistency measured whether the generated output explicitly retained the original drought probability value without modification or omission. Semi-guided prompting demonstrated a probability consistency rate of 0.00% because generated outputs produced short interpretation-like responses without explicitly repeating the original numerical drought probability values. Yield consistency measured whether the generated output retained the original estimated yield loss value without modification or omission.

**Table 3.** Quantitative evaluation of prompting strategies for agronomic report generation.

| Prompt Type | Category Consistency (%) | Probability Consistency (%) | Yield Consistency (%) | Hallucination Rate (%) |
| --- | --- | --- | --- | --- |
| Baseline | 16.67 | 61.67 | 88.33 | 11.67 |
| Semi-Guided | 33.33 | 0.00 | 23.33 | 6.67 |
| Strict | 93.33 | 33.33 | 45.00 | 18.33 |

Hallucination rate was estimated by first manually reviewing generated outputs across all prompting strategies and constructing a verified list of unsupported words and phrases directly observed in generated responses. The resulting hallucination lexicon included unsupported scientific references, unsupported environmental reasoning, and unrelated economic terminology. Generated outputs containing one or more hallucination indicators were flagged as hallucinated responses.

The quantitative results indicate that strict prompting substantially improved category consistency and structured output preservation compared to unconstrained prompting approaches. Semi-guided prompting demonstrated comparatively lower hallucination frequency due to shorter and less descriptive generated outputs, while strict prompting frequently produced structurally constrained outputs containing partially fabricated explanatory reasoning. Table 4 presents representative outputs generated using different prompting strategies.

**Table 4.**
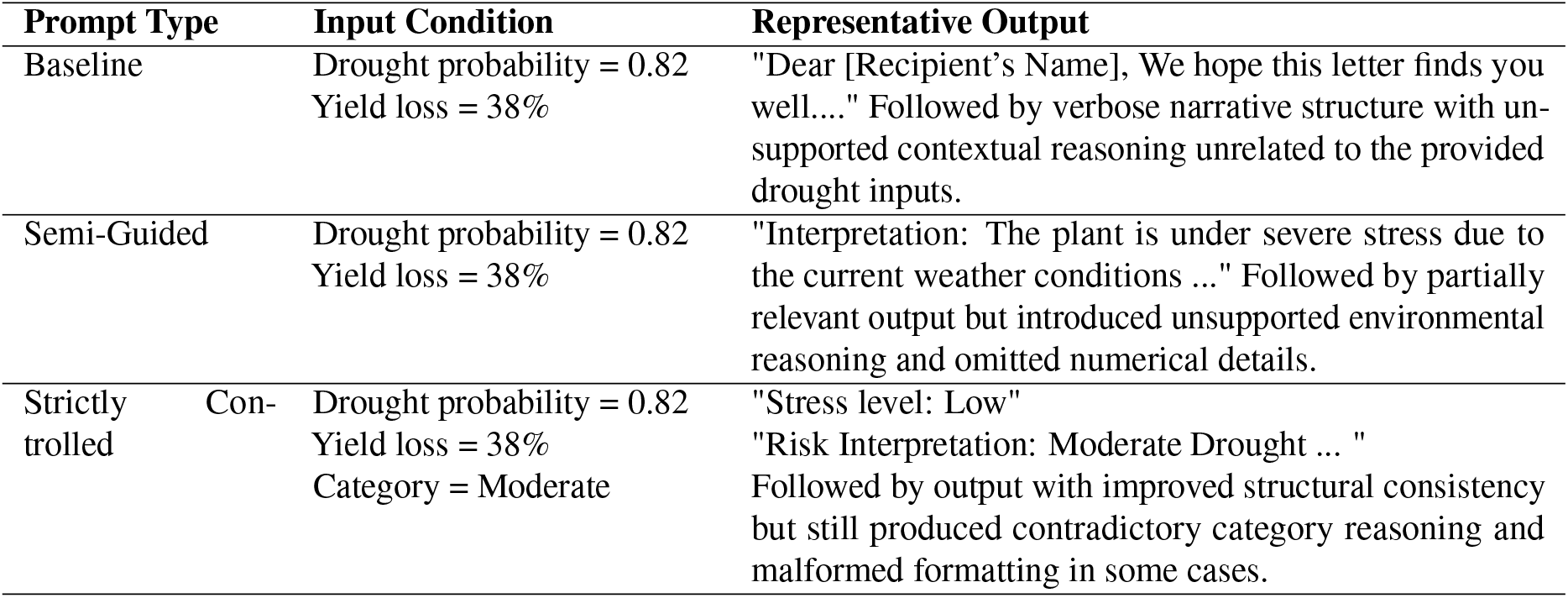
Representative outputs generated using different prompting strategies in the AgriLLaMA-based agronomic reporting framework.

### RGB-Based Visible Drought-Severity Results

All 44 selected RGB images were successfully processed, and the resulting masks contained valid canopy pixels with no unexpected class values. The mean RGB-derived drought severity index (DSI) was 0.2758 ± 0.2732, corresponding to an average visibly stressed canopy fraction of 27.58%. The median DSI was 0.14, with an interquartile range of 0.04–0.49 and an overall range of 0.0068–0.91. These values indicate substantial variation in visible canopy stress among the selected plants.

The complementary healthy fraction had a mean of 0.72 ± 0.27 and a median of 0.86. Its interquartile range was 0.5052–0.9590, with values ranging from 0.0871 to 0.9932. Fig. 8 presents the healthy-green and visibly stressed canopy fractions for individual plants ranked by increasing DSI. Fig. 9 shows the distribution of plant-level DSI values. The distribution was right-skewed, with many plants exhibiting relatively low visible stress and a smaller subset showing moderate-to-high stressed-canopy fractions. Overall, the results demonstrate that the RGB-based analysis represents visible canopy stress as a continuous plant-level trait rather than assigning each plant to a single healthy or drought-stressed category.

**Figure 8.**
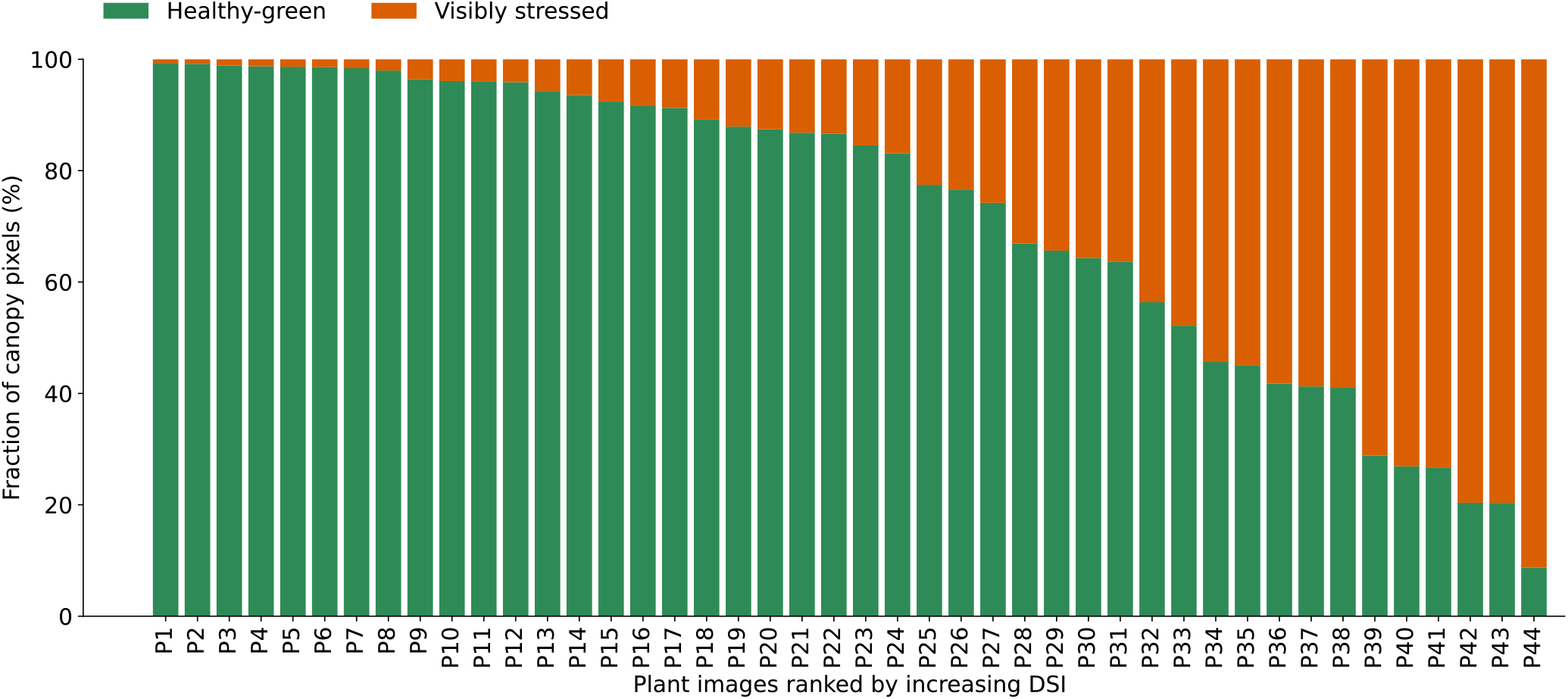
Healthy-green and visibly stressed canopy fractions for 44 individual plant images ranked by increasing RGB-derived drought severity index (DSI), with P1 representing the lowest DSI and P44 the highest.

**Figure 9.**
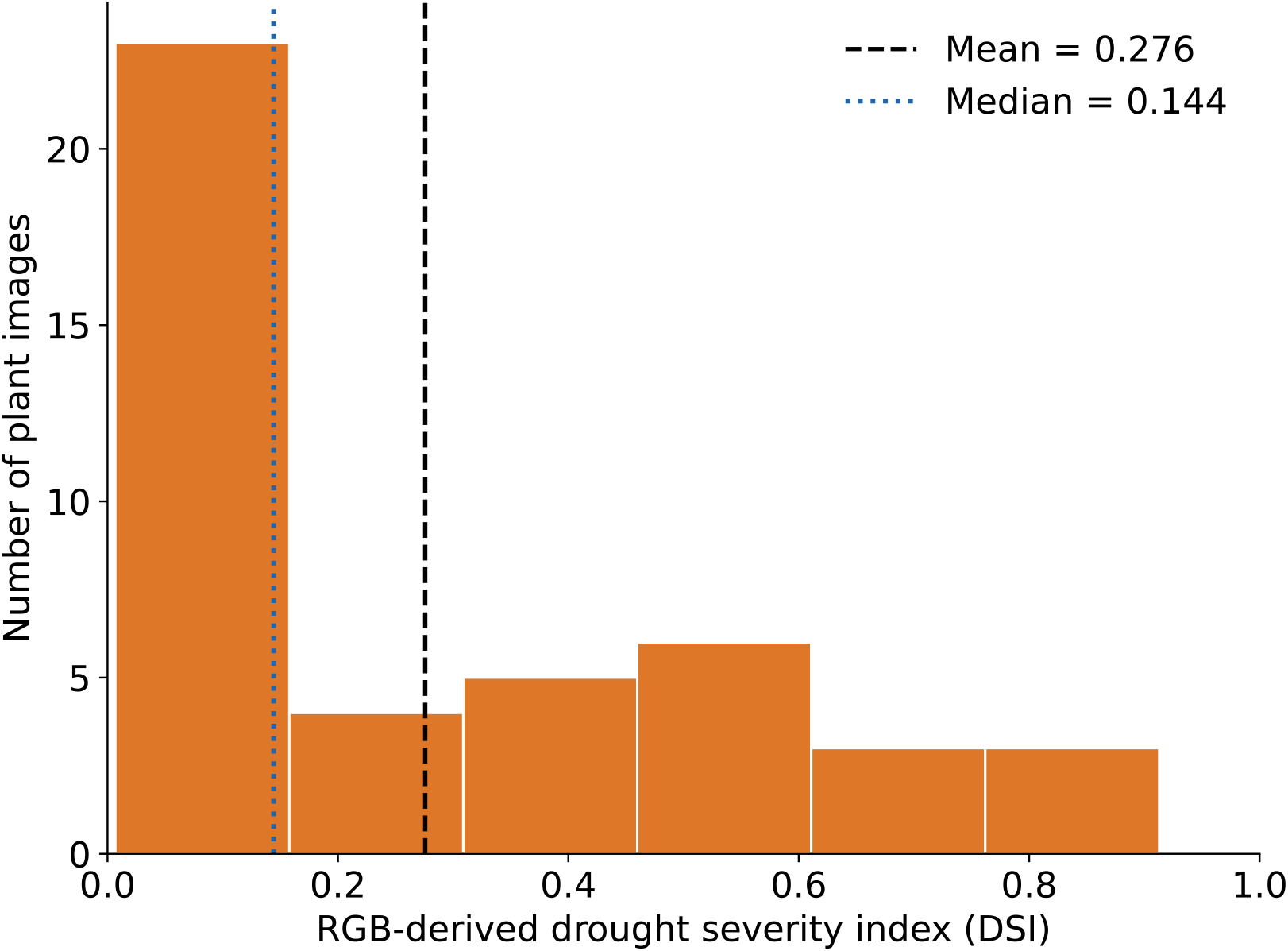
Distribution of plant-level DSI values. The dashed and dotted vertical lines indicate the mean (0.2758) and median (0.1442), respectively.

## Discussion

The proposed Drought-Spec-Net framework demonstrates strong performance for early drought stress detection using Vis–NIR spectral data. However, in the context of this study, drought classification is not the final objective but rather the foundational step in a broader decision-support pipeline. The primary goal is to transform raw spectral measurements into actionable agronomic insights through a sequential process involving stress detection, potential yield-impact mapping, and automated reporting.

On the original dataset (IR = 2), the model achieves an accuracy of 97.37%, precision of 95.24%, recall of 98.25%, and an F1-score of 96.61%, confirming its effectiveness in learning discriminative spectral representations for drought stress classification. The confusion matrix further shows that the model correctly identifies nearly all drought-stressed samples with zero false negatives, which is particularly important in agricultural applications where missed stress detection can lead to severe yield losses. In addition, the ROC-AUC of 99.72% confirms excellent separability between classes.

A comparative analysis with baseline and state-of-the-art models (Table 1) shows that the proposed method consistently outperforms all competing approaches across all evaluation metrics. Classical machine learning (RF) performs significantly worse, indicating its limitation in capturing complex spectral patterns. Among deep learning models, 1D-CNN shows competitive performance but still lags behind the proposed architecture in F1-score and recall.

State-of-the-art models such as 1D-SP-Net and 1D-ResGC-Net show comparatively lower performance than expected from published results. This may be attributed to the fact that these models were reimplemented based on architectural descriptions, as the original training code was not fully available. Differences in hyperparameter tuning, training strategy, and implementation details can significantly influence performance, particularly for attention-based or residual architectures. Despite this limitation, these models still provide strong benchmark baselines for evaluation.

To further evaluate robustness, experiments were conducted under varying imbalance ratios. At IR = 1 (balanced setting), performance remains comparable to the original dataset, indicating that resampling does not significantly improve generalization and that the original dataset already contains sufficient discriminative structure for learning. At IR = 10 (moderate imbalance), a decline in recall is observed for most baseline models, reflecting reduced sensitivity toward minority (drought) samples, whereas the proposed Drought-Spec-Net maintains relatively stable performance, demonstrating better resilience to moderate distribution shifts. At IR = 50 (severe imbalance), most baseline models degrade significantly, with several collapsing in terms of F1-score due to strong majority-class bias, although some models retain high accuracy, this is misleading under extreme imbalance conditions. In contrast, the proposed model remains comparatively more stable, although performance degradation is still observed, highlighting the inherent difficulty of learning under extreme class skew.

Beyond classification, the framework translates drought probabilities into exploratory, literature-informed potential yield-impact indicators, providing a preliminary agronomic interpretation of model outputs. Most test samples were categorized as No Loss, while smaller subsets were assigned to Low, Moderate, and Severe potential yield-impact categories. These mapped values should not be interpreted as validated yield-loss predictions because the spectral dataset does not contain measured plant-level yield data.

Finally, the integration of the AgriLLaMA-based agronomic reporting module enhances the practical usability of the system by converting numerical predictions into structured agronomic summaries. This improves interpretability and supports decision-making in precision agriculture. Despite these advantages, the AgriLLaMA model suggest that all three prompting strategies (baseline, semi-guided, and strictly controlled) exhibit some degree of uncontrolled reporting behavior. Strict prompting improves output structure. However, it does not fully eliminate hallucination or schema violations. Overall, the results indicate that the LLM-based reporting layer is useful for generating agronomic reports from model outputs, but it requires rule-based validation and post-processing before deployment.

The preliminary WVSU RGB analysis demonstrated the feasibility of quantifying visible canopy stress using a continuous DSI. Unlike the spectral model–based potential yield-loss estimates, the RGB-derived DSI was not converted into yield loss because a direct relationship between visible canopy stress and yield has not yet been established. More broadly, spectral signatures and image-derived canopy traits could quantify phenotypic dynamics between individual plants, populations, and the underlying genetic architecture across spatial and temporal scales^33^. As such, the framework presented in this study could support the discovery of previously unknown phenes. Although the spectral and RGB datasets in this study were collected independently and were not paired, they demonstrate how complementary sensing modalities can characterize different aspects of plant responses to drought. Future studies using paired RGB, spectral, physiological, and yield measurements could provide a more comprehensive understanding of drought-response dynamics and validate relationships among visible stress, physiological changes, and productivity.

## Conclusion and Future Work

This study presented Drought-Spec-Net, a hybrid 1D convolutional neural network designed for early drought stress detection in tomato plants using visible–near-infrared (Vis–NIR) spectral data. The proposed model integrates local convolutional feature extraction with global spectral representation learning, enabling robust discrimination between healthy and drought-stressed samples. Experimental results demonstrate that Drought-Spec-Net achieves strong classification performance, with an accuracy of 97%, precision of 95%, recall of 98%, and an F1-score of 97%, confirming its effectiveness for early-stage drought stress detection from spectral signatures.

Beyond classification, a literature-informed potential yield-impact mapping was used to convert Drought-Spec-Net probabilities into exploratory, scenario-based productivity-risk indicators. For the 76 test samples, the mapped potential impact values ranged from 0% to 60%, with a mean of 12.97%. Because the spectral dataset did not include measured plant-level yield, these values should not be interpreted as validated predictions of actual yield loss. Instead, they demonstrate a preliminary approach for improving the agronomic interpretation of drought-classification probabilities.

Furthermore, the integration of the AgriLLaMA-based agronomic reporting module enables automatic generation of structured and human-readable summaries from model outputs. This enhances interpretability and supports practical decision-making in precision agriculture by converting numerical predictions into actionable agronomic insights. Overall, AgriLLaMA provides an effective interface for translating model predictions into agronomically meaningful reports, facilitating end-to-end decision support from spectral analysis to management recommendations.

In a separate and independent exploratory experiment, 44 RGB images, each representing a different individual tomato plant, were analyzed to quantify healthy-green and visibly stressed canopy fractions. The analysis demonstrated the feasibility of deriving a continuous RGB-based visible canopy stress index at the plant level. Because the RGB and spectral datasets were independent and unpaired, the RGB-derived index was not used to validate Drought-Spec-Net predictions or converted into potential yield impact.

Future work will evaluate Drought-Spec-Net using multi-location, multi-season, and independent datasets and investigate cross-validation and additional imbalance-handling strategies. Future greenhouse experiments will integrate paired longitudinal RGB images, spectral measurements, physiological traits, biomass, and measured fruit yield to validate relationships among visible canopy stress, physiological response, biomass reduction, and actual yield loss. Further work will also improve the PLLaMA-based reporting module using larger domain-specific datasets, deterministic numerical verification, and rule-based safeguards to reduce hallucinations and improve factual consistency.

## Acknowledgements

The authors acknowledge the institutional support provided by West Virginia State University for conducting the greenhouse experiments.

## Author contributions

K.H. conceived and designed the study, developed the methodology and computational models, analyzed and interpreted the results, supervised the research, and wrote the original manuscript. A.M. implemented and evaluated the large language model component, drafted the corresponding sections, and reviewed and revised the manuscript. E.S., Z.W., N.C., L.M.N. and L.K. collected and preprocessed the RGB tomato plant image data. K.C. contributed to tomato plant maintenance, greenhouse operations, and experimental support. P.N. provided access to greenhouse facilities and research resources and contributed to the experimental design and study coordination. A.B. provided critical review and editorial feedback on the manuscript. All authors reviewed and approved the final manuscript.

## Data availability

The public Vis–NIR spectral dataset analyzed in this study is available at https://github.com/tariyktu/1D-SP-Net/tree/main. No new spectral data were generated in this study. The original WVSU greenhouse RGB images and their corresponding three-class masks are available at https://github.com/PRISM-Research-Lab/Tomato-Drought-Impact-Assessment.

## Code availability

The Python code used to implement Drought-Spec-Net, generate the literature-informed potential yield-impact indicators, and calculate the RGB-derived visible canopy stress index is available at https://github.com/PRISM-Research-Lab/Tomato-Drought-Impact-Assessment. The repository also contains the three-class RGB masks, instructions for accessing the public Vis–NIR spectral dataset, and a link to the ilastik mask-creation tutorial.

## Additional information

Correspondence and requests for materials should be addressed to K.H.

